# No support for the adaptive hypothesis of lagging-strand encoding in bacterial genomes

**DOI:** 10.1101/2020.01.14.906818

**Authors:** Haoxuan Liu, Jianzhi Zhang

**Affiliations:** Department of Ecology and Evolutionary Biology, University of Michigan, Ann Arbor, Michigan 48109, USA

**Author notes:** Correspondence to Jianzhi Zhang, Department of Ecology and Evolutionary Biology, University of Michigan, 4018 Biological Sciences Building, 1105 North University Avenue, Ann Arbor, MI 48109, USA. Arising from Merrikh & Merrikh *Nature Communications* http://doi.org/10.1038/s41467-018-07110-3.

## Abstract

Genes are preferentially encoded on the leading instead of the lagging strand of DNA replication in most bacterial genomes^1^. This bias likely results from selection against lagging-strand encoding, which can cause head-on collisions between DNA polymerases and RNA polymerases that induce transcriptional abortion, replication delay, and possibly mutagenesis^1^. But there are still genes encoded on the lagging strand, an observation that has been explained by a balance between deleterious mutations bringing genes from the leading to the lagging strand and purifying selection purging such mutations^2^. This mutation-selection balance hypothesis predicts that the probability that a gene is encoded on the lagging strand decreases with the detriment of its lagging-strand encoding relative to leading-strand encoding, explaining why highly expressed genes and essential genes are underrepresented on the lagging strand^3,4^. In a recent study, Merrikh and Merrikh proposed that the observed lagging-strand encoding is adaptive instead of detrimental, due to beneficial mutations brought by the potentially increased mutagenesis resulting from head-on collisions^5^. They reported empirical observations from comparative genomics that were purported to support their hypothesis^5^. Here we point out methodological flaws and errors in their analyses and logical problems of their interpretation. Our reanalysis of their data finds no evidence for the adaptive hypothesis.

---

Following Merrikh and Merrikh^5^, we refer to the leading-strand encoded genes as co-directional (CD) genes, because the movement of DNA and RNA polymerases in these genes are co-directional, and refer to lagging-strand encoded genes as head-on (HO) genes. Merrikh and Merrikh’s primary evidence for the adaptive hypothesis was their inference that the fraction of present-day HO genes that were previously CD exceeds the fraction of present-day CD genes that were previously HO in each of six bacterial species examined, which they interpreted as natural selection for HO under the reasonable assumption that the inversion mutation from HO to CD and that from CD to HO have equal rates per gene. However, if this interpretation were true, the number of HO genes would gradually rise and eventually exceed the number of CD genes, contradicting the preponderance of CD genes in most bacterial genomes. The cause of the above paradox is that, to infer selection, one should compare the CD-to-HO inversion rate with the HO-to-CD inversion rate. But the CD-to-HO rate does not equal the fraction of present-day HO genes that were CD, but the fraction of previously CD genes that are now HO. The same can be said for the HO-to-CD rate. Based on Merrikh and Merrikh’s estimates of previously and present-day HO genes and CD genes, we found that the rate of inversion from CD to HO is lower than the reverse rate in four of the six species examined (Table 1), consistent with the prediction of the mutation-selection balance hypothesis but opposite to that of the adaptive hypothesis. In fact, a lower inversion rate from CD to HO than the converse rate was reported almost 20 years ago in each of four bacterial species examined then^6^. Merrikh and Merrikh’s mistake is puzzling given that they cited this study in their paper^5^.

**Table 1.** HO-to-CD and CD-to-HO inversion rates based on the inversions inferred by Merrikh and Merrikh.

| Species | Current<br># of CD genes | Current<br># of HO genes | HO-to-CD<br>inversions | CD-to-HO<br>inversions | HO-to-CD<br>inversion rate<sup>1</sup> | CD-to-HO<br>inversion rate<sup>2</sup> | *P*-value<sup>3</sup> |
| --- | --- | --- | --- | --- | --- | --- | --- |
| *Mycoplasma gallisepticum* | 651 | 159 | 166 | 121 | 0.81 | 0.20 | <10<sup>-5</sup> |
| *Staphylococcus aureus* | 2216 | 656 | 84 | 428 | 0.27 | 0.17 | <10<sup>-5</sup> |
| *Bacillus subtilis* | 3104 | 1139 | 127 | 777 | 0.26 | 0.21 | 7.4×10<sup>-3</sup> |
| *Campylobacter jejuni* | 1048 | 651 | 23 | 529 | 0.16 | 0.34 | <10<sup>-5</sup> |
| *Klebsiella pneumonia* | 2999 | 2405 | 465 | 1102 | 0.26 | 0.30 | 2.3×10<sup>-3</sup> |
| *Mycobacterium tuberculosis* | 2415 | 1665 | 603 | 712 | 0.39 | 0.28 | <10<sup>-5</sup> |

<sup>1</sup>Number of HO-to-CD inversions divided by the number of previously HO genes, which is the number of current HO genes plus the number of HO-to-CD inversions minus the number of CD-to-HO inversions.
<sup>2</sup>Number of CD-to-HO inversions divided by the number of previously CD genes, which is the number of current CD genes plus the number of CD-to-HO inversions minus the number of HO-to-CD inversions.
<sup>3</sup>Chi-squared test of equal rates between HO-to-CD and CD-to-HO inversions.

Furthermore, Merrikh and Merrikh’s inference of previously HO genes and previously CD genes is error-prone^5^. For each genic region, they computed the leading strand GC skew by the difference in frequency between G and C, relative to the total frequency of G and C. They assumed that a negative GC skew means that the gene has been recently inverted^5^, based on a tendency for the leading strand to have a positive GC skew due to mutational bias^7^. It should be noted that the G and C frequencies of a genic region are influenced not only by mutation but also by protein-related selection. To the best of our knowledge, no study has used the GC skew alone to infer gene inversion in bacteria. Comparison of gene orientation determined from genome assemblies is the standard method to infer gene inversion in bacterial (and organelle) genomes, while the GC skew is sometimes used as confirmatory evidence^6,8,9^. Even when the GC skew is used as a confirmation, the common practice is to calculate it at third codon positions or four-fold degenerate sites, because nucleotide frequencies at these sites are subject to weaker protein-related selection^6,8,9^. By contrast, Merrikh and Merrikh computed the GC skew of a gene using its entire coding region, further increasing the likelihood of erroneous inferences of gene inversion.

We thus used the standard method to infer gene inversion in the six species analyzed by Merrikh and Merrikh. For each focal species A, a closely related species B from the same genus and an outgroup species C were selected, and orthologs among the three species were identified by reciprocal best hits from protein BLAST analysis (Supplementary Table 1). Gene orientation was determined from reference genome assemblies and inversions were inferred using the parsimony principle. We then counted the number of inversions in the lineage leading to the focal species A from the common ancestor of A and B. The results showed that the rate of CD-to-HO inversion is significantly different from the rate of HO-to-CD inversion in three of the six species. In all three cases, the former rate is lower than the latter rate (Table 2), again supporting the mutation-selection balance hypothesis but refuting the adaptive hypothesis. Under the assumption that the number of HO genes observed from a species has reached the equilibrium, the number of CD-to-HO inversions should equal the number of HO-to-CD inversions.

**Table 2.** HO-to-CD and CD-to-HO inversion rates based on the inversions inferred by the standard method.

| Species | Current<br># of CD genes | Current<br># of HO genes | HO-to-CD<br>inversions | CD-to-HO<br>inversions | HO-to-CD<br>inversion rate<sup>1</sup> | CD-to-HO<br>inversion rate<sup>2</sup> | P-value<sup>3</sup> |
| --- | --- | --- | --- | --- | --- | --- | --- |
| *Mycoplasma gallisepticum* | 271 | 48 | 11 | 21 | 0.290 | 0.075 | $3.5 \times 10^{-5}$ |
| *Staphylococcus aureus* | 1175 | 310 | 16 | 26 | 0.053 | 0.022 | $3.4 \times 10^{-3}$ |
| *Bacillus subtilis* | 1646 | 406 | 8 | 30 | 0.021 | 0.019 | 0.71 |
| *Campylobacter jejuni* | 553 | 309 | 68 | 89 | 0.236 | 0.155 | $3.6 \times 10^{-3}$ |
| *Klebsiella pneumoniae* | 1515 | 1169 | 3 | 9 | 0.0026 | 0.0059 | 0.20 |
| *Mycobacterium tuberculosis* | 930 | 450 | 8 | 22 | 0.018 | 0.023 | 0.56 |

<sup>1</sup>Number of HO-to-CD inversions divided by the number of previously HO genes, which is the number of current HO genes plus the number of HO-to-CD inversions minus the number of CD-to-HO inversions.
<sup>2</sup>Number of CD-to-HO inversions divided by the number of previously CD genes, which is the number of current CD genes plus the number of CD-to-HO inversions minus the number of HO-to-CD inversions.
<sup>3</sup>Chi-squared test of equal rates between HO-to-CD and CD-to-HO inversions.

However, we observed that the former is greater than the latter in each species (Table 2). This is likely because the evolutionary time concerned (since the divergence of species A from B) is relatively short such that a sizable fraction of deleterious CD-to-HO inversions have yet to be purged. Such a time lag in the effect of purifying selection is well known^10^. It is notable that we identified 114 HO-to-CD and 197 CD-to-HO inversions in the six species, while Merrikh and Merrikh identified an order of magnitude more inversions (1468 and 3669, respectively) in the same species^5^. Apart from the general unreliability of Merrikh and Merrikh’s method, the large disparity may also be due to the fact that we aimed to detect inversions that occurred since the divergence between species A and B, while inversions detected by Merrikh and Merrikh could have occurred at any time (though older inversions have lower detectabilities). The fact that only 1 HO-to-CD and 145 CD-to-HO inversions we detected are also detected by Merrikh and Merrikh suggests that their method not only has potential false positive errors but also makes numerous false negative errors. Regardless, as aforementioned, even the inversion rates computed from their error-prone inferences supports the mutation-selection balance hypothesis rather than the adaptive hypothesis (Table 1).

In addition to analyzing gene inversions, Merrikh and Merrikh estimated the synonymous (*d*_S_) and nonsynonymous (*d*_N_) nucleotide substitution rates of individual genes^5^. They reported that *d*_S_ is not significantly different between HO and CD genes, but *d*_N_ and *d*_N_/*d*_S_ are significantly higher for HO than CD genes. The *d*_S_ comparison suggests that the point mutation rate is not different between HO and CD genes in coding regions, consistent with experimental data^11,12^ and arguing squarely against the basis of the adaptive hypothesis that the (beneficial) mutation rate is higher for HO than CD genes. Merrikh and Merrikh interpreted the results on *d*_N_ and *d*_N_/*d*_S_ as evidence for positive selection on HO genes. However, higher *d*_N_ and *d*_N_/*d*_S_ could also result from a relaxation of purifying selection^13^. Given that highly expressed genes and essential genes are underrepresented among HO genes, relaxation of purifying selection seems a more reasonable interpretation^14^. Similarly, the observation of a larger fraction of genes with *d*_N_/*d*_S_ > 1 among HO genes than CD genes^5^ can be explained by a relaxation of purifying selection on HO genes, because Merrikh and Merrikh did not show any *d*_N_/*d*_S_ ratio that significantly exceeds 1, the criterion for establishing positive selection^13^. Merrikh and Merrikh also reported enrichment of several functional groups among HO genes relative to CD genes^5^. This non-randomness could be a byproduct of the known differences between HO and CD genes in expression level and essentiality^3,4^, so cannot be used to support the adaptive hypothesis.

In conclusion, our reanalysis of the empirical data of Merrikh and Merrikh^5^ found no evidence for the adaptive hypothesis of lagging-strand encoding in bacterial genomes. Instead, all available data collected thus far are broadly consistent with the mutation-selection balance hypothesis.

---

**Supplementary Table 1.** Species used to infer gene inversions. Phylogenetic relations of each group are based on published phylogenies.

| Groups | Species of interest (A) | Sister species (B) | Outgroup species (C) |
| --- | --- | --- | --- |
| 1 | *Mycoplasma gallisepticum* | *Mycoplasma genitalium* | *Mycoplasma penetrans* |
| 2 | *Staphylococcus aureus* | *Staphylococcus saprophyticus* | *Staphylococcus sciuri* |
| 3 | *Bacillus subtilis* | *Bacillus licheniformis* | *Bacillus cereus* |
| 4 | *Campylobacter jejuni* | *Campylobacter lari* | *Campylobacter ureolyticus* |
| 5 | *Klebsiella pneumoniae* | *Klebsiella oxytoca* | *Escherichia coli* |
| 6 | *Mycobacterium tuberculosis* | *Mycobacterium haemophilum* | *Mycobacterium leprae* |

